# RNA Secondary Structure Prediction Using a Genetic Algorithm with a Selection Method Based on Free Energy Value and Topological Index

**DOI:** 10.1101/2024.01.24.576993

**Authors:** Giorgio Benedetti

## Abstract

This paper presents a genetic algorithm designed to predict RNA secondary structures, which utilizes selection criteria based on free energy (fitness) and topological similarity. This approach represents structural information using a simple number, facilitating comparisons between foldings. The simplified graph representation identifies similarities between structures that have the same type of branches. The results demonstrate that the algorithm identifies the final secondary structure with the same level of precision as the commonly used dynamic programming, but with the advantage of producing more optimal structures with different topologies. This approach maintains high population diversity and allows for the exploration of many suboptimal structures in parallel, avoiding the possibility of getting stuck in a local minimum. This permits the investigation of not only the structure with the minimum free energy, but also of other low-energy structures with different topologies that are closer to the natural fold.

## Introduction

RNA plays many important roles in gene regulation and cellular functions beyond its role in translation of genetic information [1,2]. For instance, RNase P is a 350-nucleotide RNA molecule that participates in tRNA maturation [3], while intron self-splicing from mRNA is carried out by small nuclear ribonucleoproteins containing short RNA sequences known as U RNAs [4,5]. Additionally, ribozymes are RNA molecules that exhibit catalytic activity [6], while the Signal Recognition Particle, comprised of a 300-nucleotide RNA molecule and several proteins, is believed to influence the translocation of newly synthesized proteins by binding to ribosomes on the endoplasmic reticulum membrane [7]. RNA molecules are linear polymers of variable length that consist of an ordered sequence of nucleotides, each containing one of four nitrogenous bases: adenine (A), guanine (G), cytosine (C), and uracil (U) (primary structure).

RNA has the tendency to fold onto itself, creating intramolecular hydrogen bonds between certain base pairs, which results in the formation of a secondary structure [8]. Unlike protein folding, which is primarily driven by hydrophobic forces, the RNA folding process is hierarchical [8]. The formation of the RNA secondary structure, which consists of base pairs, occurs rapidly in linear RNA with a significant decrease in energy, whereas the development of a complex tertiary structure is typically much slower [9].

An RNA secondary structure can be viewed as a group of substructures that can be categorized into various elementary motifs, as shown in Figure 1. It contains both double-stranded (DS) and single-stranded (SS) regions. In the DS regions, the molecule folds and twists into a double helix with a minimum length of two base pairs, forming a stem.

**Fig 1.**
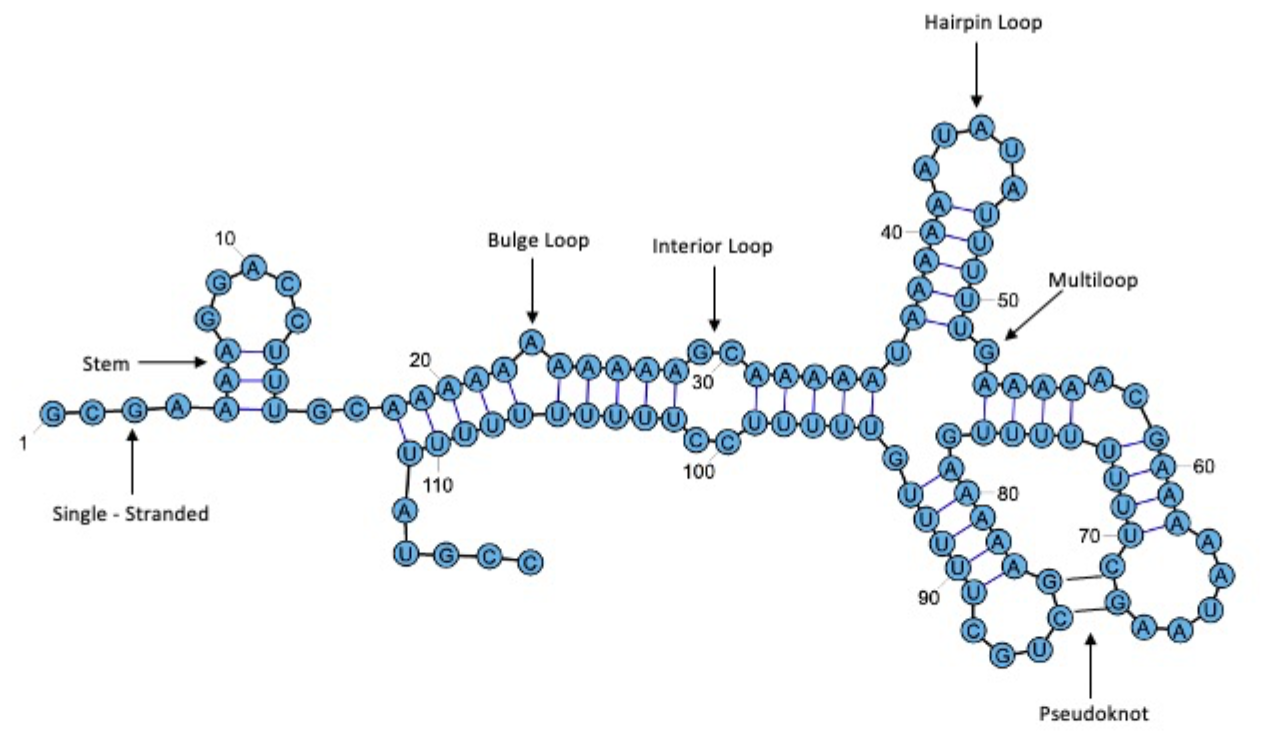
Basic structural motifs in RNA secondary structures

Many base pairings in RNA follow the classic Crick-Watson pairing rules, such as G:C or A:U, but other types of pairings exist - the G:U pairing, for example, which is known as a wobble pair (WP) [10,11]. Non-standard pairings have been observed in a variety of artificially synthesized and natural RNA molecules.

In terms of topology, the single-stranded (SS) regions can be classified into five categories: hairpins (H), which fold the strand and are believed to have a minimum single-stranded loop length of three nucleotides; bulges (B), which are unpaired nucleotides found only in one chain of the double helix; interiors (I), which consist of one or more unpaired bases on each strand of the double helix; dangling ends (D), which are unpaired terminal regions; and multiloops (M), which are the connection zones between three or more double-stranded regions.

Another crucial structural motif is the pseudoknot (PK).

In RNA secondary structure, a pseudoknot is a type of RNA structure that occurs when a single-stranded region of RNA folds back and forms base pairs with another, non-adjacent single-stranded region, resulting in a “knot-like” structure (Fig. 1). A simple pseudoknot is defined when there is a relationship between two base pairs (*i, j*) and (*k, l*), where: *i < k < j < l*, i.e., the two base pairs cross over each other in the sequence.

Pseudoknots are often found in the non-coding regions of RNA molecules, such as in ribosomal RNA and transfer RNA. They can have important roles in regulating RNA function, for example by controlling translation or RNA stability. They can also be found in the catalytic core of group I introns, RNase P RNAs, and in the mRNA-ribosome interaction during the initiation of translation and during frameshift regulation [12].

RNA secondary structure determination is a crucial area of research in bioinformatics. It is a challenging task to predict the actual pairing among all the possible bonds that will form in a true RNA structure. The number of possible structures increases exponentially with the length of the RNA sequence, prohibiting an exhaustive search for all the structures [13,14]. However, the hierarchical nature of RNA folding allows prediction of the RNA secondary structure using computational methods.

There are two methods for predicting RNA secondary structure: the *comparative approach* and the *thermodynamic approach*. Comparative sequence analysis determines conserved base pairs between homologous sequences [15-17]. The presence of a double-helix region is considered proven when an equivalent pairing is maintained in homologous RNA, despite differences in the nucleotide sequence. A major limitation of this method is that it requires a large set of homologous sequences. Currently, only thousands of RNA families are known [18], which makes it impossible to obtain homologous sequences for all RNA.

On the other hand, the thermodynamic approach is based on the search for the most thermodynamically stable structure among all possible secondary structures that an RNA sequence can achieve. The energy rules of the thermodynamic approach are based on the *nearest neighbor energy model (NNE)* proposed by Turner et al. [19-22]. This model assumes that the stability of an RNA secondary structure depends on the local sequence features near a given pair of nucleotides. The most thermodynamically stable structure is the *Minimum Free Energy (MFE)* structure.

The parameters of the model were derived from optical melting experiments, the most commonly used experimental approach to determine the change in free energy of RNA structures. The NNE model has three restrictions on the type of structural elements to which it can be applied: hairpin loops must consist of at least 3 bases; each base can only pair with another base; no presence of pseudoknots. In this approach, it is assumed that in a secondary structure, each part of the molecule in one of the elementary motifs defined above contributes additively to the total structural stability. Thermodynamically, structural stability is measured in terms of decrease in free energy obtained when the linear molecule folds into a compact structure, and lower free energy indicates a more stable structure. The molecule with the lowest free energy is considered the most probable structure in equilibrium with other forms of the same molecule. The energy contributions of adjacent base pairs (stacking) are advantageous due to their stacking on top of each other to form regular helices. The formation of loops is often, but not necessarily, unfavorable; the exact values depend on the type of the loop, the nucleotides near the loop-closing base pair, and the exact loop sequence and whether the loop nucleotides make a stable and structured motif.

### Dynamic algorithms

The thermodynamic approach is used in recursive algorithms (or dynamic programs), widely used [23-31]and available online [32-35], to predict the most likely secondary structure. The dynamic programming approach solves problems by decomposing the original problem into subproblems. The sum of free energies available from each optimal sub-structure is calculated for all possible combinations of sub-structures, and the combination with the lowest free energy is accepted as the most likely secondary structure. The observed structure may not always be the one with the lowest free energy due to external interactions, such as solvent effects. Additionally, a large number of structures can exist within a small range of free energy values, and sometimes these structures can be vary widely.

The original recursive algorithm [36] only produces a single optimal fold. However, while the lowest energy structure is typically considered to be the natural fold, this is not always the case. External factors can often impact the resulting structure, leading to observed structures that do not necessarily have the lowest free energy [37-39]. Furthermore, within a narrow range of free energy values, a multitude of structures can exist, some of which may be vastly dissimilar. This issue becomes more pronounced as the length of the sequence increases [40]. Figure 2 displays several secondary structures of tRNA^Phe^ that have very similar free energy differences but exhibit different structures. In this particular instance, the structure with the minimum energy does not correspond to the native structure.

**Fig 2.**
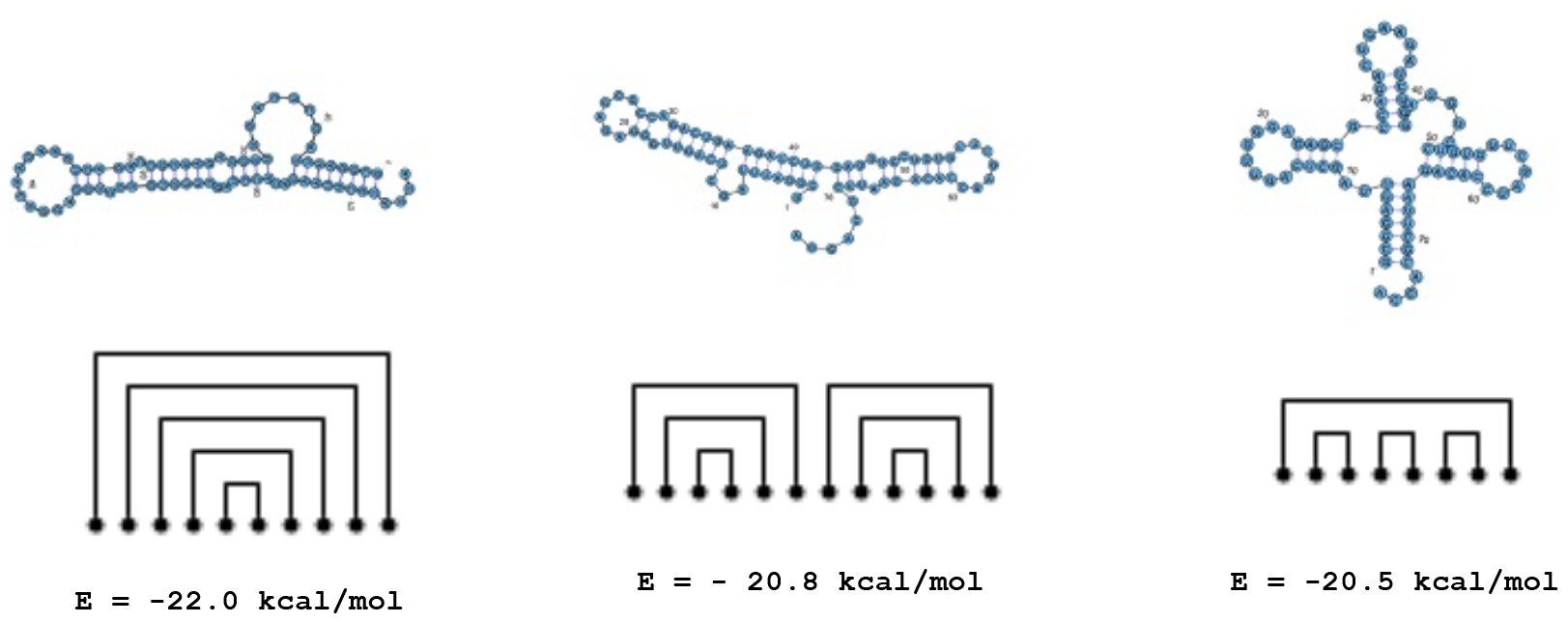
The lowest free energy secondary structure of yeast tRNA^Phe^

In addition to the inherent limitations of predicting RNA secondary structures using thermodynamic data, there are also systematic limitations due to inaccurate parameters. Some of these parameters are extrapolated, while others measured in vitro may differ from those in vivo. However, modern dynamic algorithms are capable of predicting multiple folding scenarios and incorporating experimental data such as chemical modifications and phylogenetic information, which can improve the accuracy of final structure predictions [41].

Despite these advancements, the majority of algorithms based on the minimum free energy approach do not account for the presence of pseudoknots. This is partly due to the complexity of developing such algorithms, as well as the lack of accurate thermodynamic data available for this purpose [42].

### Genetic Algorithms

A method suggested for searching RNA secondary structures is the Genetic Algorithm (GA) [43]. The Genetic Algorithm is an adaptive heuristic approach based on the concept of natural and genetic selection, which has been developed for artificial systems to achieve the robustness and adaptive properties of natural systems. Although randomized, GAs are not random but rather manipulate historical information to direct the search in the region of best performance within the search space [44].

GAs have been proven to be useful and efficient when:

1. the search space is too complex and extensive to perform an exhaustive search;
2. conventional search methods cannot provide good solutions within a reasonable time.

In general, a GA operates with several solutions (known collectively as the population) that are randomly selected. These solutions are typically encoded as binary strings. A fitness value is assigned to each solution, or individual, which is directly related to the problem’s objective function and optimization. Subsequently, the population is modified into a new population by applying three operators borrowed from genetics: *reproduction, crossover*, and *mutation*. The GA operates iteratively by successively applying these three operators to each generation until a termination criterion is satisfied [45]. The principle behind genetic algorithms is based on the concept of biological evolution. Essentially, biological evolution can be thought of as a search method that operates within a vast space of possible solutions comprised of all genetic sequences. The goal is to produce highly adapted organisms with strong survival and reproduction capabilities in a changing environment, which then transmit their genetic material to future generations. This type of algorithm is based on the Darwinian principle that the organisms best suited to their environment have the greatest chance of survival and of passing on their traits to their offspring. Thus, GAs simulates a population of individuals that evolves from generation to generation through mechanisms analogous to sexual reproduction and gene mutation. In this way, the algorithm performs a heuristic search that prioritizes the areas of the search space most likely to result in better solutions, while also exploring the less promising areas that may still yield useful results using fewer resources.

Genetic algorithms have been used across a wide range of problems, from aircraft design to modeling biological systems [46,47]. Their versatility stems from their ability to efficiently solve problems when the *building block hypothesis* is effective. The building block hypothesis suggests that a better solution can be obtained by assembling short *patterns* or *schemes* that can be recovered from some other inferior solution. The algorithm’s efficiency is due to its implicit parallelism, which allows it to manipulate many possible solutions simultaneously.

The building block hypothesis is well-suited for the additive rule used to calculate the free energy of secondary structures. Additionally, since GAs produce a population of optimal solutions, it is possible to study not only the minimum free energy structure but also other low-energy structures that may be closer to natural folding.

GAs have been applied to RNA secondary structure prediction since the early 1990s, and there have been many advances in the field [48-54]. The key elements of a GA include chromosomal representation, selection, crossover, mutation, and fitness function calculation. The GA procedure used in this work can be summarized as follows (see figure 3).

1. The first step is to search for all helices that can potentially contribute to the development of secondary structures.
2. A diverse initial population Y of *n* structures (chromosomes) is randomly selected while attempting to maintain the highest diversity of the population. The fitness (free energy) is then calculated for each structure.
3. P1 and P2 chromosomes are selected from population Y based on their fitness value and topological characteristics.
4. The crossover operator is applied to P1 and P2 to create offspring O1 and O2.
5. The offspring chromosomes are then added to the initial population, and a mutation operator is applied to the entire population.
6. Only solutions with the best fitness are kept from the new generation Y’, which replaces the old population.
7. The same procedure is repeated on the new population obtained, and the iteration is terminated when the value of the fitness of the best structure (elite structure) does not change and remains constant a parameter such as average fitness, Hamming distance, structure degeneration, etc.

**Fig 3.**
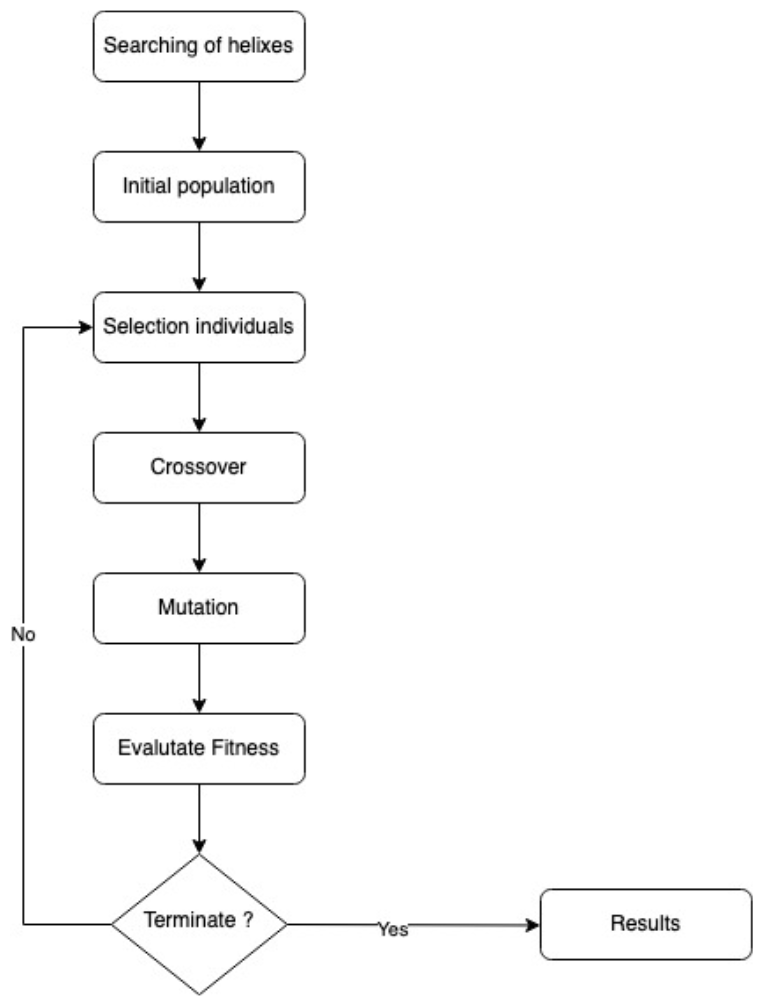
Genetic algorithm process

In this version of GA, significant importance was given to step 3, which aimed to maintain a high level of genetic variability in the population to avoid obtaining suboptimal solutions and RNA secondary structures that differ widely from natural ones. To achieve this, the selection process considered not only the free energy but also the topology of the individuals within the population. The following section outlines the primary steps and operators that make up the specific genetic algorithm used to search for RNA secondary structures.

### Search for all helices obtainable from an RNA folding

The first step of the algorithm involves searching for all possible helical regions that can be formed by the RNA sequence. In a previous study [55], a novel approach based on the convolution theorem was developed and is outlined here. Each helix can be visualized as the result of folding the sequence around a specific nucleotide. This is equivalent to comparing the sequence with its reverse complement and shifting it appropriately (as shown in figure 4). If f(x) represents a discrete function, its auto-convolution [56] is given by equation (1), where u represents the nucleotide position in the sequence and x denotes the shift of the sequence:

**Fig 4.**
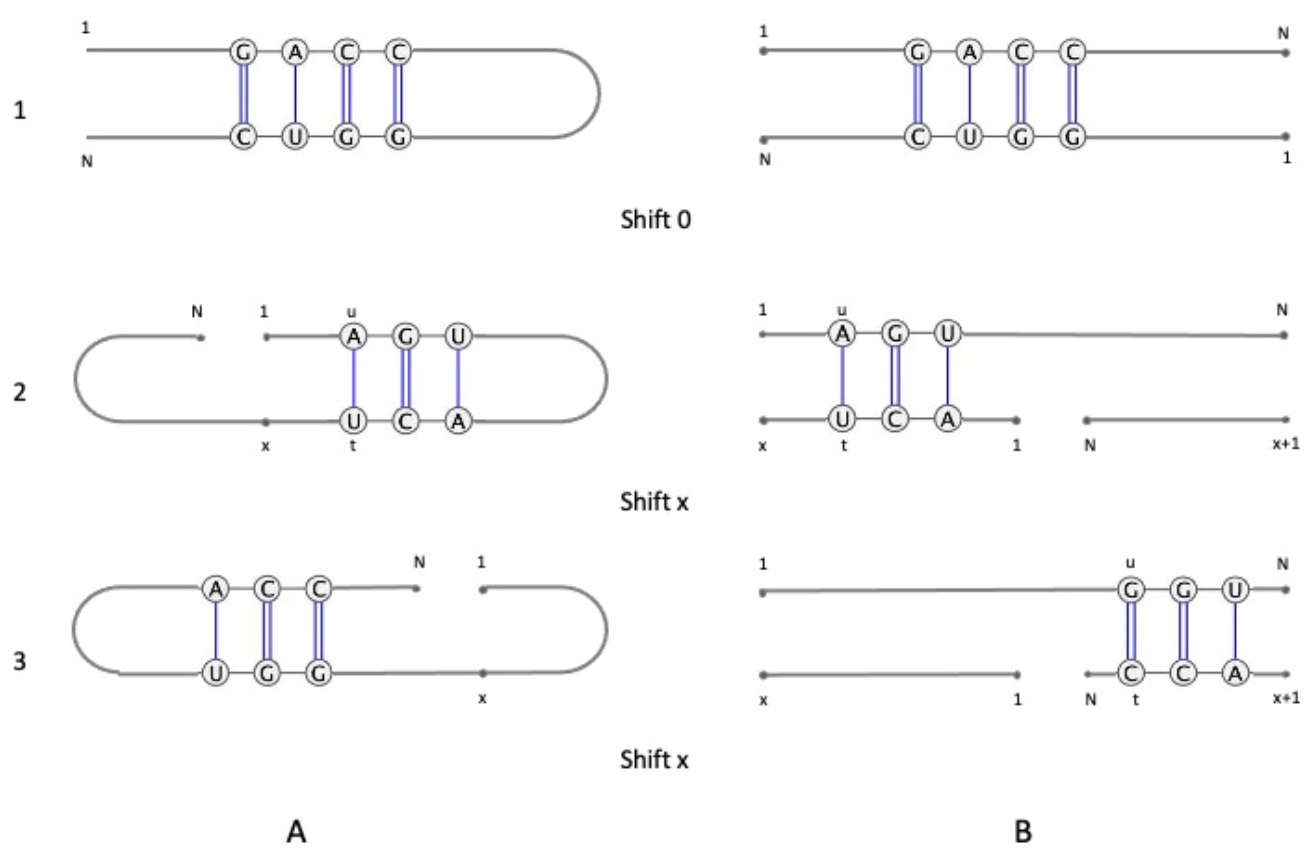
A) Physical folding of the sequence around a nucleotide; B) the corresponding folding obtained by shifting the sequence along itself reversed. 1) shift x = 0; 2) generic x shift

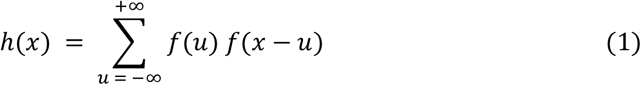

For an RNA sequence, the values of u and x that are significant lie between 1 and N, where N represents the total number of nucleotides in the sequence. Consequently, equation (1) can be expressed as:

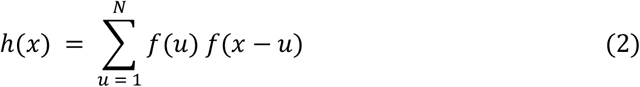

Every shift represents a fold of the sequence, and doing the auto-convolution of the nucleotide chain involves observing the relationship between the bases facing each other (i.e., whether or not they pair, as shown in figure 4b). *f(u)* is defined such that the product *f(u)f(x-u) ≠ 0* only when the corresponding nucleotides are paired by hydrogen bonds. A tetra-dimensional complex vector is then assigned to each base (as seen in table I) so that the dot product is nonzero only in cases of Crick Watson (G:C, A:U) or Wobble (G:U) pairing. The value of *h(x)* represents the number of paired nucleotides for each fold. By expanding the size of the vectors assigned to the bases, other non-standard pairings can also be considered. In this algorithm, the Turner energy tables were used, and only standard pairings were considered. The bases coded as S in table 1 indicate that they are modified in a way that prevents them from forming Crick Watson pairings.

**Table 1.** Tetradimensional complex vector assignments

| Table 1 – Tetradimensional complex vector assignments |  |
| --- | --- |
| | $f(u)$ |
| G | $(2, 2i, 2, 2i) * 1/\sqrt{8}$ |
| A | $(2, 2i, -2, -2i) * 1/\sqrt{8}$ |
| U | $(2, -2i, 0, 0) * 1/\sqrt{8}$ |
| C | $(1, -i, 1, -i) * 1/\sqrt{8}$ |
| S | $(0, 0, 0, 0)$ |

Assignment to find Crick-Watson and Wobble pairing. Bases coded as S are blocked by modifications at key groups and they cannot form hydrogen bonds

**Table 2.** Comparison of the structures with lowest energy obtained with the genetic algorithm and the RNAStructure program in terms of STY, PPV, and F-m.

| RNA sequence | Length | n structures | PK | Genetic Algorithm |  |  |  |  |  | RNAstructure |  |  |  |  |  |
| --- | --- | --- | --- | --- | --- | --- | --- | --- | --- | --- | --- | --- | --- | --- | --- |
|  |  |  |  | No Shape |  |  | Shape |  |  | No Shape |  |  | Shape |  |  |
|  |  |  |  | STY | PPV | F<sub>m</sub> | STY | PPV | F<sub>m</sub> | STY | PPV | F<sub>m</sub> | STY | PPV | F<sub>m</sub> |
| Pre-Q1 riboswitch, B. Subtilis | 34 | 50 | 1 | 62,5 | 100,0 | 76,9 | 62,5 | 100,0 | 76,9 | 62,5 | 100,0 | 76,9 | 62,5 | 100,0 | 76,9 |
| Fluoride riboswitch, P. syringae | 66 | 100 | 1 | 62,5 | 90,9 | 74,1 | 62,5 | 100,0 | 76,9 | 56,3 | 64,3 | 60,0 | 62,5 | 71,4 | 66,7 |
| Adenine Riboswitch | 71 | 100 | 0 | 100,0 | 100,0 | 100,0 | 100,0 | 100,0 | 100,0 | 100,0 | 100,0 | 100,0 | 100,0 | 100,0 | 100,0 |
| tRNA Phe | 76 | 100 | 0 | 95,7 | 100,0 | 97,8 | 100,0 | 100,0 | 100,0 | 95,2 | 100,0 | 97,5 | 100,0 | 100,0 | 100,0 |
| TPP riboswitch | 79 | 100 | 1 | 77,3 | 85,0 | 81,0 | 95,5 | 87,5 | 91,3 | 77,3 | 85,0 | 81,0 | 95,5 | 87,5 | 91,3 |
| SARS corona virus pseudoknot | 82 | 200 | 1 | 65,4 | 68,0 | 66,7 | 65,4 | 68,0 | 66,7 | 69,2 | 90,0 | 78,2 | 69,2 | 75,0 | 72,0 |
| Cyclic di-GMP riboswitch | 97 | 200 | 0 | 75,0 | 77,0 | 76,0 | 89,3 | 86,2 | 87,7 | 75,0 | 77,0 | 76,0 | 89,3 | 86,2 | 87,7 |
| SAM 1 riboswitch | 118 | 100 | 1 | 74,4 | 80,6 | 77,4 | 76,9 | 85,7 | 81,1 | 74,4 | 80,6 | 77,4 | 76,9 | 85,7 | 81,1 |
| 5S rRNA | 120 | 250 | 0 | 28,6 | 25,0 | 26,7 | 97,1 | 91,9 | 94,4 | 28,6 | 25,0 | 26,7 | 85,7 | 76,9 | 81,1 |
| M-Box riboswitch, B. subtilis | 154 | 300 | 0 | 87,5 | 91,3 | 89,4 | 87,5 | 91,3 | 89,4 | 87,5 | 91,3 | 89,4 | 87,5 | 91,3 | 89,4 |
| P546 domain, b13 group I intron | 155 | 300 | 0 | 80,4 | 81,8 | 81,1 | 96,4 | 96,4 | 96,4 | 42,9 | 44,4 | 43,6 | 94,6 | 96,4 | 95,5 |
| Lysine riboswitch, T. maritima | 174 | 400 | 1 | 81,0 | 89,5 | 85,0 | 81,0 | 89,5 | 85,0 | 77,8 | 83,1 | 80,3 | 79,4 | 89,3 | 84,0 |
| Group I intron, Azoarcus sp | 214 | 500 | 1 | 66,7 | 71,2 | 68,9 | 76,2 | 80,0 | 78,0 | 76,2 | 75,4 | 75,8 | 81,0 | 85,0 | 83,0 |
| Signal recognition particle RNA, human | 301 | 600 | 0 | 22,0 | 24,7 | 23,3 | 59,0 | 59,6 | 59,3 | 0,0 | 0,0 | 0,0 | 59,0 | 59,0 | 59,0 |
| Hepatitis C virus IRES domain | 336 | 600 | 1 | 45,2 | 45,6 | 45,4 | 79,8 | 86,5 | 83,0 | 39,4 | 38,0 | 38,7 | 79,8 | 86,5 | 83,0 |
| RNAse P, B. Subtilis | 401 | 800 | 1 | 43,4 | 45,7 | 44,5 | 64,4 | 69,2 | 66,7 | 57,4 | 55,0 | 56,2 | 76,5 | 81,5 | 78,9 |
| Group II intron, O. iheyensis | 412 | 900 | 1 | 90,2 | 99,2 | 94,4 | 87,9 | 98,3 | 92,8 | 88,6 | 97,5 | 92,8 | 74,3 | 84,5 | 79,1 |
| Group I intron, T. thermophila | 425 | 900 | 1 | 90,1 | 84,4 | 87,2 | 90,1 | 85,6 | 87,8 | 83,2 | 74,3 | 78,5 | 87,8 | 88,6 | 88,2 |
| 5 domain of 16S rRNA, H. volcanii | 473 | 1000 | 0 | 84,0 | 75,2 | 79,3 | 88,2 | 83,0 | 85,5 | 79,9 | 71,4 | 75,4 | 89,6 | 82,7 | 86,0 |
| 5' domain of 16S rRNA, E. coli | 530 | 1100 | 0 | 61,5 | 53,2 | 57,0 | 89,9 | 80,6 | 85,0 | 61,5 | 56,9 | 59,1 | 89,9 | 80,6 | 85,0 |
| 5' domain of 23S rRNA, E. coli | 511 | 1100 | 0 | 69,8 | 55,0 | 61,5 | 92,4 | 75,3 | 83,0 | 92,4 | 76,4 | 83,7 | 93,3 | 74,5 | 82,8 |
| Average |  |  |  | 69,7 | 73,5 | 71,1 | 83,0 | 86,4 | 84,1 | 67,9 | 70,7 | 68,9 | 82,6 | 84,9 | 83,4 |

The table presents the STY, PPV, and F-m values obtained for the set of examined sequences. The first three columns show the corresponding sequence lengths, the number of structures used in the initial population, and the number of pseudoknots in the native structure. The following columns report the values obtained with and without SHAPE data, from both the proposed genetic algorithm and the RNAstructure program. The *Scorer* program, which is available as a part of the RNAstructure package, was used to calculate the STY and PPV values by comparing the predicted secondary structure to an established structure [77].

In this version of the search algorithm, only structures that do not result in pseudoknots - structures in which there are only relationships of inclusion and exclusion between two helices - are considered. This type of structural motif is not considered during the analysis of secondary structures, but can be searched for in an “optimization” phase of the obtained foldings.

In the initial phase of the algorithm, the stacking energies for each identified helix are calculated using the parameters of the proposed NNE energy models. Additionally, the contribution of experimental data can be included as a pseudo-energy (see below) without significantly increasing the computation time. The topological relationships between each helix and all others are used to generate an incompatibility matrix. This matrix is then utilized in the search algorithm to compare the topological relations between the various helices.

### Encoding Scheme

A chromosome is represented by a secondary structure without knots and stored as a set of helices. For a given RNA sequence, if there are N possible helices, the chromosome can be represented as a string of N bits (Fig. 5a). Each bit represents the presence (1) or absence (0) of a particular helix. This is a low-cardinality alphabet representation that is more efficient in genetic algorithms [57]. However, the structure is actually stored as a set of helices to reduce memory usage (Fig. 5b).

**Fig 5.**
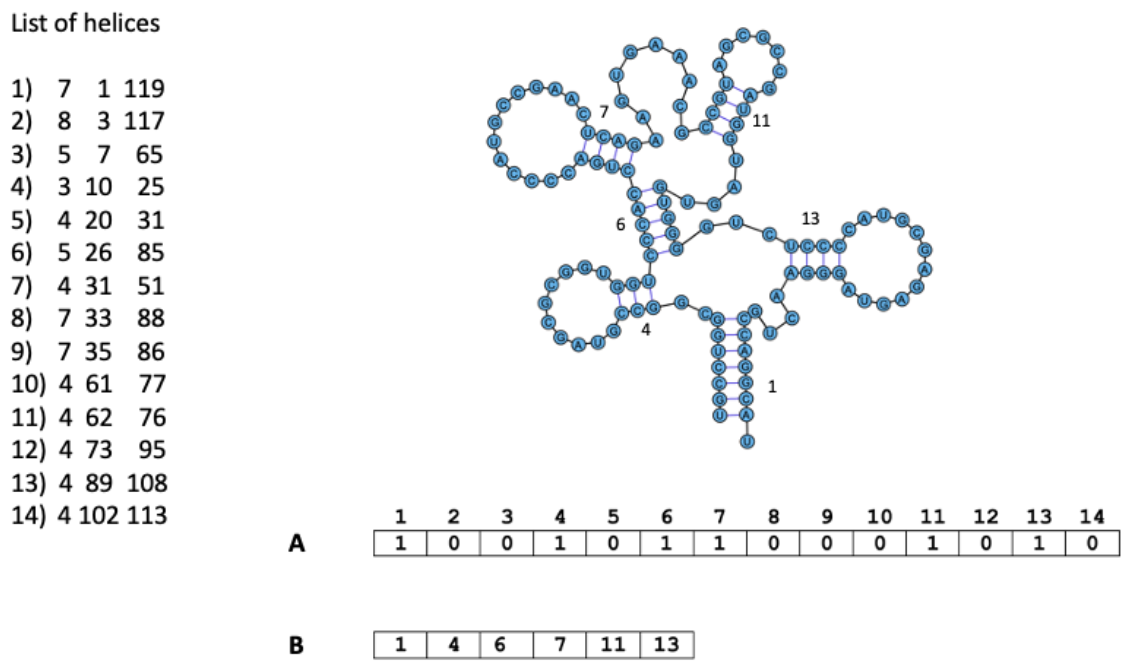
Secondary structure and its record representation. Numbers 1 to 14 indicate the helices in an ordered list. The numbers in the list indicate respectively: order number of helix; number of base-pairs forming helix; first nucleotide of the 5’ strand of helix, in the sequence numbered from 5’ to 3’ end; final nucleotide of 3’ strand of helix. In record A the bit position in position *i* indicates the presence (1) or absence (0) of helix *i*. Record B is compressed representation actually used in the algorithm.

### Diversity and selective pressure

Diversity and selective pressure are crucial for genetic algorithms to avoid premature convergence of solutions. In order for evolution to occur, solutions with higher fitness must have a higher probability of surviving and reproducing, but this selection can lead to favoring a sub-optimal solution at the expense of all competitors. An excessive number of identical individuals in a population can cause evolution stagnation, as crossover between two identical solutions can only produce a clone. If not properly programmed, GAs can lead to stagnation after a few generations due to the limited size of potential solutions populations. Once a solution that is better than others arises, it tends to prevail over all others, reducing the chances of further improvement [58].

To ensure that the solution space is adequately searched - especially in the initial optimization phases - it is necessary to maintain a diverse population of solutions [59]. Larger populations lead to better outcomes due to the larger pool of different schemas available [60]. However, premature convergence occurs when a GA population reaches a sub-optimal state, making genetic operators unable to produce offspring that improve upon their parents. Therefore, it is necessary to maintain a balance between genetic diversity and selective pressure to ensure that the GA converges to the optimal global solution in a reasonable time [61].

The algorithm seeks to balance these two factors by acting on the selection of the initial structure pool and the criterion for choosing individuals that gives rise to crossover. Furthermore, elitism and insertion of new random structures during iteration execution are introduced to increase population diversity. Elitism involves keeping one or more optimal solutions obtained in a generation, called *elites*, in the next generation without undergoing any changes. This strategy speeds up the convergence of the algorithm as it avoids the loss of the best-found structure. The injection of new structures allows for the recovery of some lost helices from previous iterations.

#### 1. Initial population selection

The population size for this algorithm depends on the number of possible helices, which in turn is determined by the length and base composition of the sequence. To create an initial population with a diverse set of helices, helices are randomly chosen using the roulette method [62], where helices with better energy have a higher probability of being selected. The selected helix is added to the individual in training only if it does not create nodes with helices already present. The number of attempts to add new helices is fixed, but the number of helices in each individual can vary. Only structures with negative free energy are accepted. The helices with the lowest energy are selected (about 15% of the total) and their presence in the population is limited. The approach is to form the most stable helices first and then add the less stable ones. In this way one can limit the number of initial structures to a reasonable number with a significant variety of helices. Other helices will gradually be taken into account during the iteration process by the mutation operators.

#### 2. Selection of parent structures for crossover

During the crossover phase (CO) (described later), the initial structures are randomly selected to form the mating pool. Then, pairs of structures that generate CO are chosen based on two criteria: a) *quality rank*, which is determined by fitness, and b) *diversity rank*, which is based on the topological diversity of the structures [63]. This process is repeated until the desired number of individuals to be selected from the mating pool (percentage of CO) is achieved.

#### a) Quality rank: fitness function

The fitness function provides an estimation of an individual’s quality and determines its probability of reproduction. It is calculated based on the change in free energy resulting from the formation of a secondary structure, which is evaluated using energy tables that rely on the NNE model.

#### b) Diversity rank: topological diversity of structures

The second criterion for selecting parent structures that generate crossover is based on the concept of ecological niche [64]. A niche is defined as a subset of the environment that can support different types of life. Within a niche, physical resources are limited and must be shared among the population. Niching methods are often employed to naturally create niches and species within the search space. These methods help to maintain population diversity and allow the genetic algorithm to explore multiple suboptimal structures in parallel, preventing it from getting stuck in a local minimum of the search space [65].

In the algorithm, the selection of two structures that will undergo crossover is favored by selecting the most topologically similar structures, which are believed to belong to the same niche. Here, “similar” refers to structures with the same number of branches, regardless of their branch lengths.

This approach suggests representing secondary structures as graphs similar to molecular graphs, where the loops are the vertices and the helices are the edges (Fig. 6a and 6b). To account for differences in branch length, vertices connecting only two edges are excluded, resulting in a simplified graphical representation (Fig. 6c). To compare these structural graphs, a *topological index* is used. Topological indices are calculated by applying mathematical rules to graphs in order to obtain numbers that can correlate properties with gradual changes in structures, effectively discriminating between isomers and accounting for dimensional dependencies [66].

**Figure 6.**
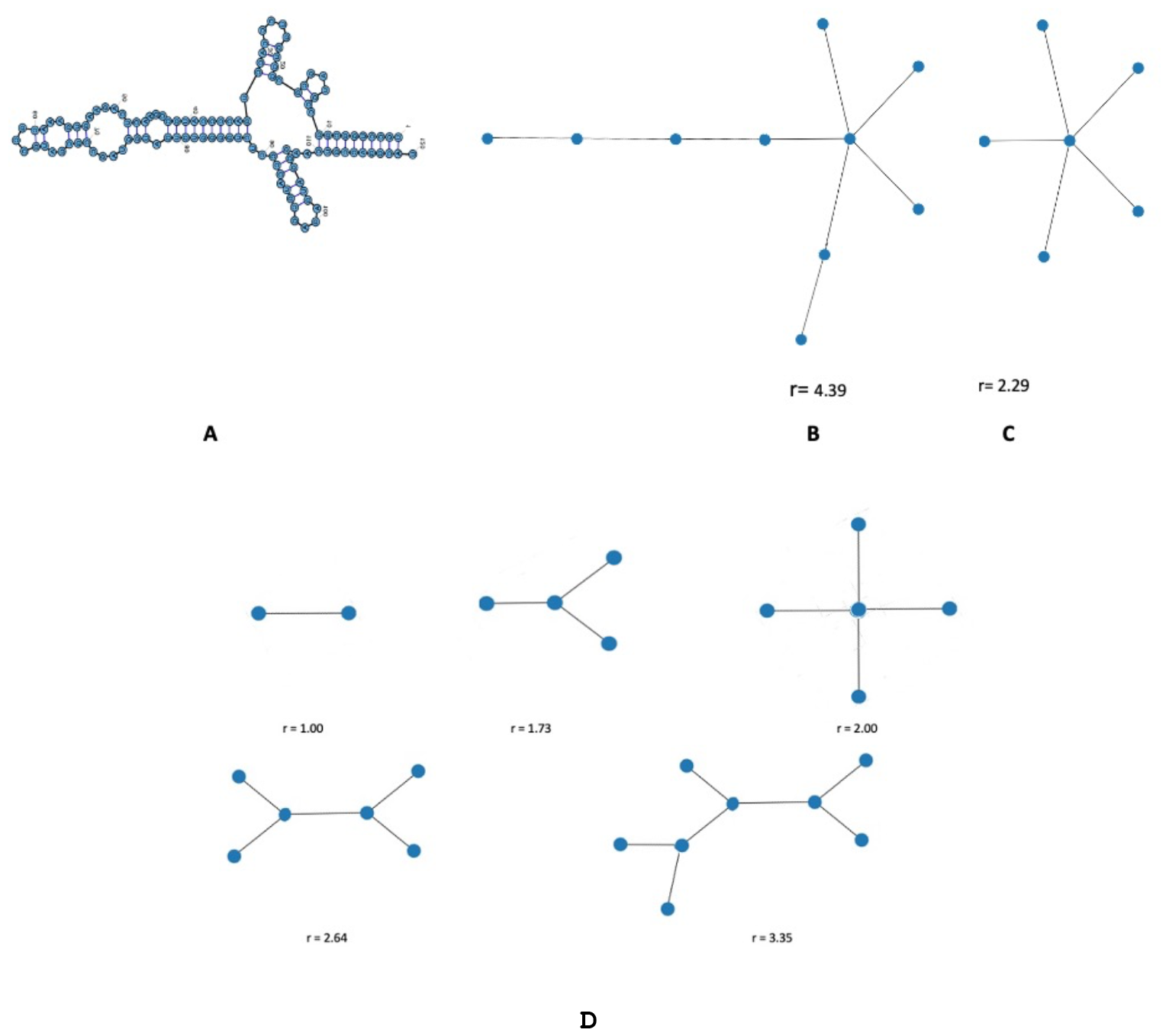
RNA secondary structure, its graphical representation, and the Randić topological index. (A) shows the putative suboptimal free energy secondary structure. (B) presents the structural graph representation with vertices representing the loops and edges representing the helices. (C) is a simplified version of the structural graph representation with omitted vertices that connect only two edges. (D) demonstrates some of the graphical representations in increasing order of complexity, along with the reported Randić topological indices r.

**Fig 7.**
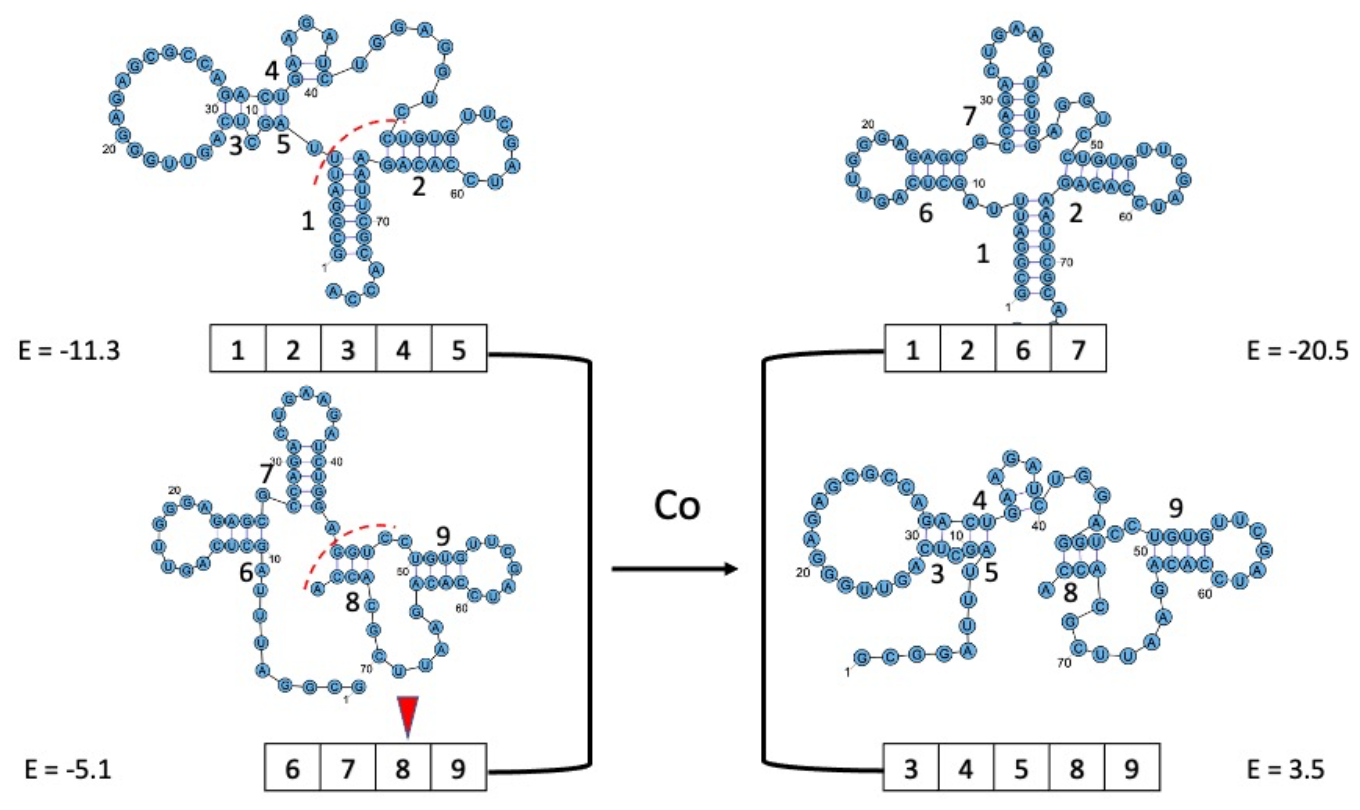
Schematic representation of reproduction with crossover: selection of two individuals (on the left) and application of the crossover mechanism to generate two offspring structures (on the right)

Among the different topological indices available, the *Randić index* was chosen based on two key criteria: the uniqueness of the index value corresponding to the graph and the ease of calculation. The Randić index is based on the topological concept of vertex degree, which is simply the number of edges connected to a given vertex. The Randić index of a graph is defined by the following formula:

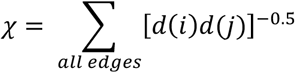

where the degree of a vertex is the number of edges incident to it, and *d(i)* and *d(j)* are the degrees of vertices *i* and *j* that define the edge *ij*. The Randić indices of various structural graphs are illustrated in Fig. 6B, 6C, and 6D. To compare two secondary structures, their simplified structural graphs are compared by checking the identity of their Randić indices. During the selection of structure pairs for CO, one structure is chosen randomly, and the other is chosen from the remaining structures that have the most similar Randić index and best energy. This selection process in the algorithm aims to obtain as many different structures as possible, rather than local variations of the same individual.

### Crossover operator

The crossover phase is the core of the genetic algorithm and is highly effective in optimizing problems where solutions can be conceptualized as blocks that contribute independently to the fitness function, which is precisely the building block hypothesis. In the crossover process, new solutions are created by combining parts from both parental solutions. This results in the generation of two new structures by exchanging structure parts from the two parent folds.

During the crossover process, two new offspring individuals are generated by exchanging bits between the two selected parent individuals. The exchange occurs at a random crossover point, and all the bits following this point are swapped between the individuals. This mechanism is appropriate for RNA secondary structure optimization as the free energy is additive. In the algorithm, a helix is randomly selected, and all the helices included in it are chosen. Then, the helices that need to be moved in the second individual to make room for the helices from the first individual are determined, and the two groups of helices are exchanged. This process involves dividing each individual into two substructures, exchanging them between the individuals, and then assembling the resulting substructures to obtain optimal and suboptimal structures. Sometimes, individuals with positive free energy may be generated during the crossover process. In such cases, a randomly chosen helix is selected, and a substitution is attempted to bring the free energy to a negative value without changing the topology. If this is not possible, the structure is replaced with a new one (see Section on selection of initial structures). Crossover can be applied to a fraction of the total structures present in the mate pool.

### Mutation

Mutation is an important operator in genetic algorithms that helps maintain genetic diversity in populations and prevents the loss of important patterns. In the RNA secondary structure prediction problem, mutation is used to randomly change the structure of a fraction of the structures in the population. This can involve adding a new helix, removing an existing one, or replacing an existing helix with a randomly chosen new one. The mutation operator is especially important in this application because it helps prevent the loss of smaller helices that may contribute moderately to the stabilization of structures, but are important for refining the stable ones in the final phases of the search.

The mutation operator is applied to a fraction of the structures in the population, typically ranging from 5% to 10%. During the course of the algorithm, an increasing number of helices are considered for the mutation operator.

### Iteration Termination

The termination of the iteration is not solely dependent on the constancy of the fitness of the best structure, but also on a selected parameter that indicates when the optimization process has reached a stopping point. This parameter can be a measure of population diversity, such as the average Hamming distance, or a measure of population degeneration, which is the similarity of secondary structures forming the population. When this parameter remains constant for a certain number of iterations, the algorithm is terminated, and the best structures are returned as the final solutions.

#### a) Average Hamming Distance

The Hamming distance [67] is a measure of the difference between two vectors, computed as the sum of the corresponding elements that differ between them. In practice, a greater Hamming distance indicates a larger difference between the vectors, while a smaller Hamming distance indicates greater similarity.

Mathematically, the Hamming distance can be expressed as follows:

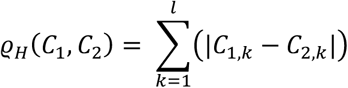

In this genetic algorithm, the average Hamming distance is used, which is calculated as the average distance between pairs of vectors in a population of N individuals. If the distance between each pair of vectors is 0, it indicates that the arrays are identical.

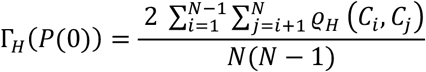

#### b) Degeneration of the population

The similarity between two structures can be assessed by counting the number of overlapping helices they have [68]. This count can be normalized and averaged over all pairs in the population to provide a measure of population degeneration.

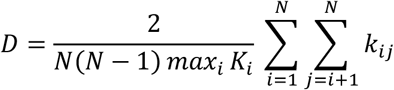

where *K*_*i*_ is the number of helices in structure *i, k*_*ij*_ is the number of helices common to structures *i* and *j*; *N* is the size of the population. *D* is equal to 1 if and only if the entire population is represented by copies of a single individual. The value of *D* ranges from 0 to 1, where *D* = 1 indicates that the entire population is represented by copies of a single individual. As the population evolves, the *D* parameter increases and tends to 1 and the calculations are terminated when the *D* value exceeds a specified threshold, D_c_.

### Including Experimental Data in the Fitness Function

Thermodynamic parameters are useful for predicting the stability of individual helices and hairpins. However, predicting large RNA structures can be difficult due to several factors. As mentioned before, many different structures can exist within a small energy range, and the NNE model’s simplifications and incomplete thermodynamic rules introduce uncertainties into fitness calculations. To improve the prediction of functional secondary structures, experimental data obtained from techniques such as SHAPE can be included (Selective 2’-Hydroxyl Acylation by Primer Extension) [69] or structure-sensitive enzymatic cleavage and chemical probing reagents.

The SHAPE technique is based on the fact that the 2-hydroxyl in the RNA ribose’s nucleophilic reactivity is sensitive to local nucleotide flexibility. SHAPE reagents react more readily with conformationally unconstrained or flexible nucleotides found in single-stranded, loop, or bulge regions, while nucleotides in structured regions are less reactive. SHAPE reactivity can be converted into pseudo-energy terms, ΔG_SHAPE_, as there is an inverse correlation with the nucleotide’s probability of forming a base pair. By adding this pseudo-energetic term to the free energies of the NNE model, the search for RNA secondary structures can be refined [70].The pseudo-energy term for each paired base *i* is calculated as follows:

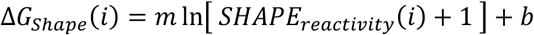

The *m* and *b* parameters in the equation above are used to scale the contribution of experimental data to the energy function. The intercept *b* represents the contribution of pseudo-energy of the nucleotide pair whose SHAPE reactivity is zero, while *m* represents the penalty assigned to nucleotides with high SHAPE reactivity. This algorithm use the values *m* = 1.8 kcal/mol and *b* = -0.6 kcal/mol, which are the default parameters in the RNAStructure program used for comparison.

Chemical probing reagents are useful for investigating the secondary structure of RNA, as they interact with RNA in different ways depending on its structure. These reagents can modify nucleotides exposed in single-stranded regions or paired, or interact with specific nucleotides that form particular secondary structures, such as hydrogen bonds. One commonly used chemical reagent for investigating RNA secondary structure is dimethyl sulfate (DMS). DMS methylates accessible guanines in single-stranded regions, which can then be detected by enzymatic digestion and electrophoretic analysis. Paired guanines are not accessible to DMS, making it possible to determine the location of single-stranded and paired regions in RNA.

Another example of a reagent used to investigate RNA secondary structure is the enzyme RNase V1. This enzyme specifically hydrolyzes RNA at the level of hydrogen bonds in paired regions, allowing the mapping of such regions through electrophoretic analysis.

In a previous study, a method for incorporating experimental data was presented. This method is summarized here and can be utilized to integrate the data into the fitness function [71].

When introducing experimental data into the fitness function, it is important to note that the data may not always be in agreement with the formation of the most energetically favorable secondary structures. For this reason, a method of minimizing the free energy of soft constraints was preferred over a prior modification of the list of double-helix regions as hard constraints, which would exclude helices in conflict with experimental data or impose energetically disfavored ones. By introducing a coefficient, the “experimental” term can be given a growing weight, gradually forcing the system to obtain families of structures with increasingly greater respect for experimental data and reaching a compromise between respecting these data and minimizing free energy.

Indicating with T the function to be minimized:

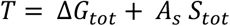

The free energy of the RNA structure is denoted as *ΔG*_*tot*_; *A*_*s*_ is a coefficient used to balance the experimental term with the energetic term; and *S*_*tot*_ is the experimental contribution. *S*_*tot*_ is determined by assigning values (*s*) to individual bases depending on the type of experimental data and normalizing them to a maximum of |*s*|=1. For instance, +1 is assigned to bases in a single strand, and -1 to bases exhibiting evidence of a double helix. If there is a broader range of values, the range of 0 to 1 can be utilized. *S*_*tot*_ can be split into two terms: *S*_*btot*_ (double-stranded regions) and *S*_*ltot*_ (single-stranded regions):

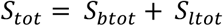

The contribution of experimental data can be represented by the equation [71]:

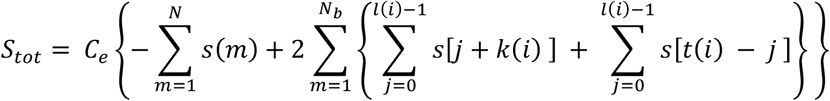

where *N* is the number of nucleotides in the sequence, *N*_*b*_ is the total number of helical regions in the structure; and *l(i), k(i)* and *t(i)* are respectively the length, starting position and final position of the helical region i. *C*_*e*_ is the reference value corresponding to the average free energy stacking value of the considered sequence.

For prediction evaluation, metrics commonly used to evaluate RNA secondary structure predictions were considered: positive predictive value (PPV), sensitivity (STY), and F-measure (F-m) [72]. PPV is a metric used to evaluate the accuracy of RNA secondary structure predictions. It represents the proportion of predicted base pairs that are true positives, i.e., the base pairs that are present in the actual RNA secondary structure.

The formula for PPV is:

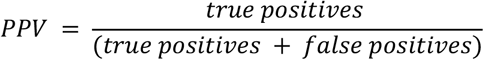

where true positives are the number of predicted base pairs that are also present in the known or experimentally determined structure, and false positives are the number of predicted base pairs that are not present in the known or experimentally determined structure.

The STY indicates how well the algorithm is able to detect the true positive base pairs, which are the base pairs that are present in both the predicted and the actual RNA secondary structures.

The formula for sensitivity is:

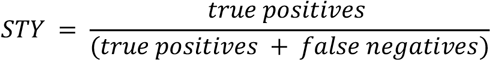

where true positives are the number of correctly predicted base pairs, and false negatives are the number of base pairs that are present in the actual structure but are not predicted by the algorithm.

The F-measure provides a more comprehensive evaluation of an algorithm’s performance. It is a combination of both STY and PVV, and provides a balanced measure of the algorithm’s accuracy.

The formula for F-measure is:

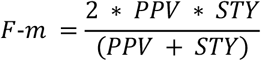

The impact of the experimental term on the total energy can be demonstrated by plotting the percentage of satisfied experimental data against the values of the experimental coefficients used. To illustrate secondary structure prediction with the addition of experimental data, an example is shown using the 5S rRNA sequence from Escherichia coli, which was selected for its abundance in experimental enzymatic, chemical, and SHAPE data [73]. Figure 8A displays the average percentage of satisfied experimental data for a set of 100 structures as a function of changes in the experimental coefficient. It can be observed that as the program is “forced” to search, it increasingly tends to select families of structures that on average increasingly satisfy the experimental data. Once a certain value is reached, further increases in experimental coefficients do not produce significant changes (Figure 8B). These pseudo-energy values can be calculated at the outset and integrated in the initial helix search phase.

**Fig 8.**
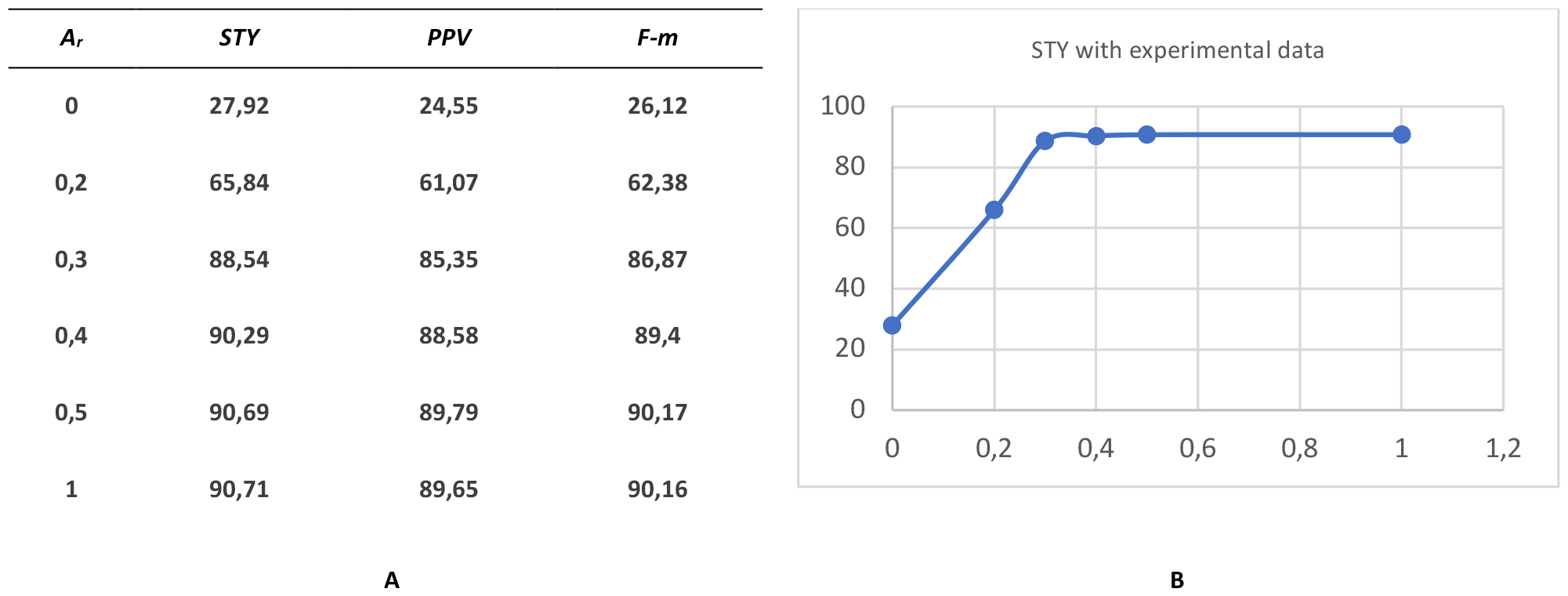
Average % values of satisfied experimental data foro a set of 100 structures as a function of variations in the experimental coefficient.

Figure 9B displays the secondary structure prediction in the absence of experimental constraints, which exhibits low values of STY, PPV, and F-m (STY = 28.57, PPV = 25.0, and F-m = 94.4). Figure 9C illustrates the secondary structure prediction with the inclusion of experimental data, which is considerably more accurate with a STY of 94.3% and a PPV of 91.7%. These values were obtained from five program runs with different initial populations consisting of 100 structures and with an experimental coefficient *A*_*s*_ = 1.0. Figure 9D presents structure prediction made using SHAPE data with the parameters of *m* = 1.8 and *b* = -0.6. In this case, the prediction exhibits high values of STY, PPV, and F-m (STY = 97.1, PPV = 91.9, and F-m = 94.4).

**Fig 9.**
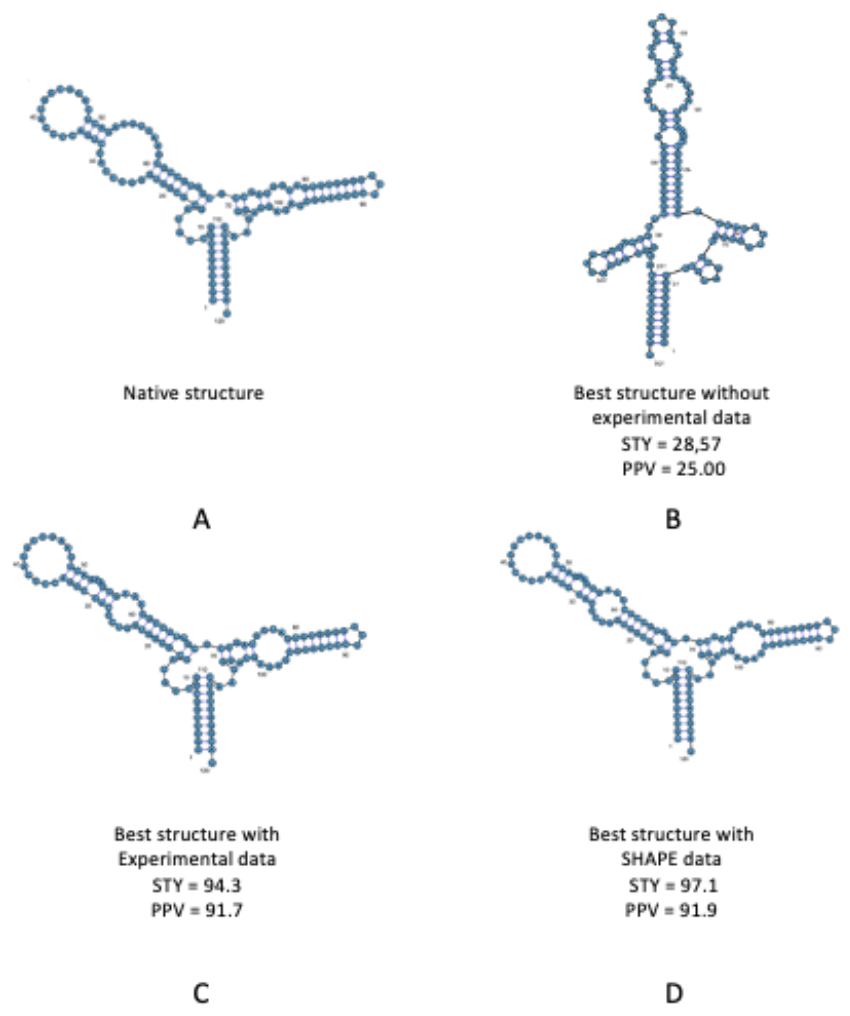
Native secondary structure of Escherichia coli 5S rRNA sequence (A). Structure predicted in the absence of experimental constraints (B), with the introduction of experimental constraints (C) and with SHAPE data (D).

### Drawing image

The computer program RiboSketch [74] was used to draw parts of the figures.

### Results

To test this genetic algorithm, it was applied to a benchmark with known reference structures [75]. This test set contains sequences, corresponding SHAPE data, and reference structures derived from X-ray crystallography experiments or predicted through comparative sequence analysis. In particular, the results were compared with RNAStructure, one of the most used dynamic programs available on the Web. The analysis was conducted on a different number of starting structures, depending on the length of the sequence. A number of starting structures equal to double the length of the sequence under analysis were taken. The number of iterations also varied greatly because the length of the RNA increased as the number of strands considered increased. In the algorithm, 600 was considered the maximum distance between base pairs because 99% of the paired bases in an RNA molecule involve pairings at a distance smaller than this value [76]. Only the structures with the lowest energy among those predicted were considered because even if more accurate structures are obtained with higher folding energy, there is no general way to identify them as better. The final results were compared with those obtained from the RNAStructure program and are reported in Table 3 and show substantial equivalence with those obtained from the RNAStructure program. The values indicated in the table refer to the best structure obtained by the algorithm. Often, the algorithm finds structures with energy that differs little from the best structure but are closer to the native structure. The results often lead to variations from known structure for two prior-stated reasons. First, the thermodynamical data used in the calculations always has some degree of imprecision. Second, the energy model used excludes pseudoknot structures, which are present in about half of the reference RNAs. The use of SHAPE-based constraints leads to better prediction results for many RNAs. In these cases, the interactions of pseudoknots are reflected in the SHAPE data themselves.

Indeed, if the predicted structure is compared with the native structure without pseudoknots, the results obtained for the optimal secondary structure are even better.

### Conclusions

The genetic algorithm used in this study to predict RNA secondary structures employs selection criteria based on both the free energy value (fitness) and the topological similarity using a topological index, such as Randić. This approach allows for structural information to be represented as a number, making it easy to compare different folding structures. The topological index is calculated based on a simplified representation of the graph, allowing for recognition of similarity between structures with the same type of branches, despite differences in length or number of helices between individual branches.

In comparison to one of the most commonly used dynamic programs, the final results show that the algorithm can identify the final secondary structure with nearly the same degree of accuracy, with a further advantage of producing a variety of optimal structures with different topologies. This approach maintains high diversity in the population and allows the GA to explore many suboptimal structures in parallel, preventing it from getting trapped in a local minimum in the search space. Consequently, the algorithm can investigate not only the structure with the lowest free energy but also other low-energy structures with different topologies, which may be closer to the natural fold.

## Acknowledgements

Thanks to Dr. Aaron Greenberg for his great availability and useful discussion

## Notes

### Competing Interest Statement

The authors have declared no competing interest.

## References

[1] Higgs PG, Lehman N. The RNA World: molecular cooperation at the origins of life. Nat. Rev. Genet. 2015; 16(1):7–17

[2] Mortimer SA, Kidwell MA, Doudna JA. Insights into RNA structure and function from genome-wide studies. Nat. Rev. Genet. 2014; 15(7):469–79

[3] Pace, N. R., Brown J. W. (1995). Evolutionary perspective on the structure and function ribonuclease P, a ribozyme. J. Bacteriol. 177, 1919–1928.

[4] Baserga S. J., Steitz, J. A. (1993). The diverse world of small ribonucleoproteins. In: The RNA World (eds. Gesteland, R. F. & Atkins, J. F.), pp. 359–381. Cold Spring Harbor Laboratory Press.

[5] Zweib C. (1997). The uRNA database. Nucleic Acids Res. 25, 102–103.

[6] Doudna J, Cech T: The natural chemical repertoire of natural ribozymes. Nature 2002, 418: 222–228. 10.1038/418222a

[7] Zweib, C., Samuelsson T. (2000). Signal recognition particle database. Nucleic Acids Res. 28, 171–172

[8] Tinoco I, Bustamante C. How RNA folds. J. Mol. Biol. 1999;293(2):271–81.

[9] Celander DW, Cech TR. Visualizing the higher order folding of a catalytic RNA molecule. Science. 1991; 251(4992):401–7

[10] Higgs, P.G., 2000. RNA secondary structure: physical and computational aspects. Q. Rev. Biophys. 33 (3) 199–253.

[11] Schuster, P., Stadler, P.F., Renner, A., 1997. RNA structures and folding: from conventional to new issues in structure predictions. Curr. Opin. Struct. Biol. 7 (2), 229–235.

[12] Staples, David W., and Butcher Samuel E. (2005). Pseudoknots: RNA Structures with Diverse Functions. PLoS Biol, 3(6), e213, 0956–0959.

[13] M. Zuker and D. Sankoff, RNA secondary structures and their prediction, Math. Biosci., vol. 46, pp. 591–621, 1984.

[14] Waterman M, Introduction to computational biology. Maps, sequences and genomes. Chapman & Hall, London (1995)

[15] Woese C, Pace N: The RNA World, chap. Probing RNA structure, function, and history by comparative analysis Cold Spring Harbor Laboratory Press, Cold Spring Harbor, NY 1993, 91–117.

[16] Gutell RR, Lee JC, Cannone JJ. The accuracy of ribosomal RNA comparative structure models. Curr. Opin. Struct. Biol. 2002; 12(3):301–10

[17] Madison JT, Everett GA, Kung H. Nucleotide Sequence of a Yeast Tyrosine Transfer RNA. Science. 1966; 153(3735):531–4.

[18] Burge SW, Daub J, Eberhardt R, Tate J, Barquist L, Nawrocki EP, et al. Rfam 11.0: 10 years of RNA families. Nucleic Acids Res. 2013

[19] Turner, D. H. & Mathews, D. H. (2009). NNDB: The nearest neighbor parameter database for predicting stability of nucleic acid secondary structure. Nucleic Acids Res. 38, D280–D282.

[20] https://rna.urmc.rochester.edu/NNDB/turner04/index.html

[21] Mathews D, Sabina J, Zuker M, Turner D (1999) Expanded sequence dependence of thermodynamic parameters improves prediction of RNA secondary structure. J. Mol. Biol. 288:911–940

[22] Xia T, Santa Lucia J, Burkard M, Kierzek R, Schroeder S, Jiao X, Cox C, Turner D (1998) Thermodynamic parameters for an expanded nearest-neighbor model for formation of RNA duplexes with Watson-Crick base pairs. Biochemistry 37:14719–14735

[23] M. Zuker, On Finding All Suboptimal Foldings of an RNA Molecule, Science, vol. 244, pp. 48–52, 1989.

[24] M. Zuker, Mfold Web Server for Nucleic Acid Folding and Hybridization Prediction, Nucleic Acids Res., vol. 31, pp. 3406–3415, 2003.

[25] M. Waterman and T. Smith, Rapid Dynamic Programming Algorithms for RNA Secondary Structure, Advances in Applied Math., vol. 7, no. 4, pp. 455–464, 1986.

[26] I.L. Hofacker, Vienna RNA Secondary Structure Server, Nucleic Acids Res., vol. 31, pp. 3429–3431, 2003

[27] R. Nussinov and A.B. Jacobson, Fast Algorithm for Predicting the Secondary Structure of Single-Stranded RNA, Proc. Natl. Acad. Sci. USA, vol. 77, no. 11, pp. 6309–6313, 1980.

[28] M. Zuker and P. Stiegler, Optimal Computer Folding of Large RNA Sequences Using Thermodynamics and Auxiliary Information,” Nucleic Acids Res., vol. 9, no. 1, pp. 133–148, 1981.

[29] T. Akutsu, Dynamic Programming Algorithms for RNA Secondary Structure Prediction with Pseudoknots, Discrete Applied Math., vol. 104, pp. 45–62, 2000.

[30] M.S. Waterman, T.F. Smith, RNA secondary structure: a complete mathematical analysis Math. Biosci., 41 (1978), pp. 257–266

[31] M. Zuker, P. Stiegler, Optimal computer folding of large RNA sequences using thermodynamics and auxiliary information, Nucleic Acids Res. 1981 Jan 10; 9(1): 133–148.

[32] RNAStructure - https://rna.urmc.rochester.edu/RNAstructureWeb/

[33] Sfold - http://sfold.wadsworth.org/

[34] UNAFold Web Server - http://www.mfold.org/

[35] Vienna RNA Package - http://www.tbi.univie.ac.at/RNA/

[36] Nussinov R, Jacobson AB., Fast algorithm for predicting the secondary structure of single-stranded RNA. Proc. Natl. Acad. Sci. U S A. 1980; 77(11):6309–13.

[37] John A. Jaeger, Douglas H. Turner, and Michael Zuker, Improved predictions of secondary structures for RNA. Biochemistry, 86:7706–7710, October 1989.

[38] Michael Zuker, John A. Jaeger, and Douglas H. Turner., A comparison of optimal and suboptimal RNA secondary structures predicted by free energy minimization with structures determined by phylogenetic comparison. Nucleic Acids Res., 19(10):2707–2714, 1991.

[39] D. H. Mathews, T. C. Andre, J. Kim, D. H. Turner, and M. Zuker., An updated recursive algorithm for RNA secondary structure prediction with improved free energy parameters. In N. B. Leontis and J. SantaLucia Jr., editors, American Chemical Society, 682, chapter 15, pages 246–257. American Chemical Society, Washington, DC, 1998.

[40] Schroeder SJ. Advances in RNA structure prediction from sequence: new tools for generating hypotheses about viral RNA structure-function relationships. J Virol. 2009 Jul;83(13):6326–34.

[41] Mathews D, Disney M, Childs J, Schroeder S, Zuker M, Turner D., Incorporating chemical modification constraints into a dynamic programming algorithm for prediction of RNA secondary structure. Proc. Natl. Acad. Sci. USA 101:7287–7292, (2004)

[42] Gultyaev, A., van Batenburg, A., and Pleij, C., An Approximation of loop free energy values of RNA H-pseudoknots. RNA, 5. 609–617 (1999)

[43] Goldberg D.E. (1989), Genetic Algorithm in search, optimization and machine learning.

[44] Sivaraj, R. and Ravichandran, T. (2011) A Review of Selection Methods in Genetic Algorithm. International Journal of Engineering Science and Technology, 3, 3792–3797.

[45] Katoch S., Chauhan S. S., and Kumar V. (2021). A review on genetic algorithm: past, present, and future. Multimedia Tools and Applications, 80(5), 8091–8126.

[46] L. Davies (Editor), Handbook of Genetic Algorithms. Van Nostrand-Reinhold, New York, 1991.

[47] R.K. Belew and L.B. Booker (Editors). Proceedings of the Fourth International Conference on Genetic Algorithms, Morgan Kaufmann, San Mateo, CA. 1991.

[48] F. H. D. van Batenburg, A. P. Gultyaev, and C. W. A. Pleij. The computer simulation of RNA folding pathways using a genetic algorithm. J. Mol. Biol., 250:37–51, 1995.

[49] F. H. D. van Batenburg, Alexander P. Gultyaev, and Cornelis W. A. Pleij. An APL-programmed genetic algorithm for the prediction of RNA secondary structure. Journal of Theoretical Biology, 174:269–280, 1995.

[50] B. A. Shapiro and J. Navetta. A massively-parallel genetic algorithm for RNA secondary structure prediction. Journal of Supercomputing, 8:195–207, 1994

[51] G. Benedetti and S. Morosetti. A genetic algorithm to search for optimal and suboptimal RNA secondary structures. Biophysical Chemistry, 55:253–259, 1995.

[52] Alexander P. Gultyaev, F. H. D. van Batenburg, and Cornelis W. A. Pleij., The computer-simulation of RNA folding pathways using a genetic algorithm. J.Mol. Biol., 250:37–51, 1995.

[53] Shapiro, B.A., Navetta, J. A massively parallel genetic algorithm for RNA secondary structure prediction. J Supercomput 8, 195–207 (1994)

[54] B. A. Shapiro, J. C. Wu, D. Bengali, and M. J. Potts. The massively parallel genetic algorithm for RNA folding: MIMD implementation and population variation. Bioinformatics, 17:137–148, 2001.

[55] Benedetti, G., De Santis, P. and Morosetti, S. (1990) J. Biomol. Struct. Dyn. 7, 1269–1277

[56] Bracewell, R. (1965), The Fourier Transform and Its Applications. McGraw-Hill. New York. pp. 24–50

[57] Sivanandam SN, Deepa SN (2008), Introduction to genetic algorithm, 1st edn. Springer-Verlag, Berlin Heidelberg

[58] Gupta, Deepti and Shabina Ghafir., An Overview of methods maintaining Diversity in Genetic Algorithms. (2012)

[59] L. Darrell Whitley. An overview of evolutionary algorithms: practical issues and common pitfalls. Inf. Softw. Technol. 43(14): 817–831 (2001)

[60] D.E. Goldberg, Genetic Algorithms in Search, Optimization and Machine Learning, Addison Wesley, Reading, MA, 1989.

[61] SAS/IML Users guide 9.3 vol 1 chapter 20 page 501–505.

[62] Jebari K (2013) Selection methods for genetic algorithms. Abdelmalek Essaâdi University. International Journal of Emerging Sciences 3(4):333–344

[63] Chaiwat J., Prabhas C., Self-adaptation mechanism to control the diversity of the population in Genetic Algorithm, International Journal of Computer Science & Information Technology (IJCSIT) Vol 3, No 4, August 2011

[64] K. A. DeJong, “An analysis of the behavior of a class of genetic adaptative systems,” Ph.D. dissertation, Univ. of Michigan, Ann Arbor, 1975.

[65] Bruno Sareni, Laurent Krähenbühl. Fitness sharing and niching methods revisited. IEEE Transactions on Evolutionary Computation, 1998, 2 (3), pp.97–106.

[66] Z. Mihalić and N. Trinajstić, A Graph–Theoretical Approach to Structure–Property Relationships, J. Chem. Educ. 1992, 69, 701–712.

[67] Pedro A. Diaz-Gomez and Dean F. Hougen. Initial Population for Genetic Algorithms: A Metric Approach. In proceeding of the International Conference on Genetic and Evolutionary Methods, Las Vegas, USA. 2007; 25–28.

[68] I. I. Titovà, D. G. Vorobiev, V. A. Ivanisenko, and N. A. Kolchanov. A fast genetic algorithm for RNA secondary structure analysis. In Russian Chemical Bulletin, International Edition, Vol. 51, No. 7, pp. 1135—1144, July, 2002

[69] Sloma MF, Mathews DH. Improving RNA secondary structure prediction with structure mapping data. Methods Enzymol. 2015; 553:91–114.

[70] Deigan KE, Li TW, Mathews DH, Weeks KM. Accurate SHAPE-directed RNA structure determination. Proc. Natl. Acad. Sci. U S A. 2009 Jan 6;106(1):97–102.

[71] Benedetti G, De Santis P, Morosetti S. A new method to find a set of energetically optimal RNA secondary structures. Nucleic Acids Res. 1989 Jul 11;17(13):5149–61

[72] Mathews DH. How to benchmark RNA secondary structure prediction accuracy. Methods. 2019;162-163:60–67

[73] Mathews DH, Disney MD, Childs JL, Schroeder SJ, Zuker M, Turner DH. Incorporating chemical modification constraints into a dynamic programming algorithm for prediction of RNA secondary structure. Proc. Natl. Acad. Sci. U S A. 2004 May 11;101(19):7287–92.

[74] https://rnastructure.cancer.gov/ribosketch

[75] Hajdin, C. E., Bellaousov, S., Huggins, W., Leonard, C. W., Mathews, D. H., and Weeks, K. M. (2013). Accurate SHAPE-directed RNA secondary structure modeling, including pseudoknots. Proc. Natl. Acad. Sci., 110(14), 5498–5503

[76] Doshi KJ, Cannone JJ, Cobaugh CW, Gutell RR (2004) Evaluation of the suitability of free-energy minimization using nearest-neighbor energy parameters for RNA secondary structure prediction. BMC Bioinformatics 5:105.

[77] Reuter JS, Mathews DH, RNAStructure: software for RNA secondary structure prediction and analysis, BMC Bioinformatics 11 (2010) 129.

